# Minor deviations from randomness have huge repercussions on the functional structuring of sequence space

**DOI:** 10.1101/706119

**Authors:** Laura Weidmann, Tjeerd Dijkstra, Oliver Kohlbacher, Andrei N. Lupas

## Abstract

Approaches based on molecular evolution have organized natural proteins into a hierarchy of families, superfamilies, and folds, which are often pictured as islands in a great sea of unrealized and generally non-functional polypeptides. In contrast, approaches based on information theory have substantiated a mostly random scatter of natural proteins in global sequence space. We evaluate these opposing views by analyzing fragments of a given length derived from either a natural dataset or different random models. For this, we compile distances in sequence space between fragments within each dataset and compare the resulting distance distributions between sets. Even for 100-mers, more than 95% of distances can be accounted for by a random sequence model that incorporates the natural amino acid frequency of proteins. When further accounting for the specific residue composition of the respective fragments, which would include biophysical constraints of protein folding, more than 99% of all distances can be modeled. Thus, while the local space surrounding a protein is almost entirely shaped by common descent, the global distribution of proteins in sequence space is close to random, only constrained by divergent evolution through the requirement that all intermediates connecting two forms in evolution must be functional.

**Significance Statement:** When generating new proteins by evolution or design, can the entire sequence space be used, or do viable sequences mainly occur only in some areas of this space? As a result of divergent evolution, natural proteins mostly form families that occupy local areas of sequence space, suggesting the latter. Theoretical work however indicates that these local areas are highly diffuse and do not dramatically affect the statistics of sequence distribution, such that natural proteins can be considered to effectively cover global space randomly, though extremely sparsely. By comparing the distance distribution of natural sequences to that of various random models, we find that they are indeed distributed largely randomly, provided that the amino acid composition of natural proteins is respected.

---

**N**atural proteins form the backbone of the complicated biochemical network that has given rise to the great variety of life on Earth. This highly interwoven framework of reactions seems impossible to have arisen by chance, simply because the great majority of random protein sequences fails to form a specific structure, let alone possess chemical activity. Features that distinguish naturally evolved from random sequences are therefore of great interest, both in order to understand protein evolution (1, 2) and to guide the design of new proteins (3, 4).

Searches for such differences have hitherto focused on the exhaustive enumeration of short peptides and their statistical analysis by exact occurrence (5, 6). These studies showed that the natural frequency of most peptides is similar to that expected from random sequences with the same composition. Nevertheless, the frequency of some peptides was found to deviate substantially from random occurrence, an observation which was variously discussed in terms of homologous descent and convergence due to structural and functional constraints. This enumeration approach quickly reaches its limits at sequence lengths above five, due to the fact that there are simply not enough natural sequences to populate the exponentially growing sequence space. Furthermore, pentapeptides are far from having a relevant length for understanding protein sequences. Even if proteins are dissected into their constituent domains, relevant sequence lengths still mostly range above 80 residues (7). At a complexity of 20^80^, it is clear that this sequence space cannot be analyzed by an enumeration approach.

Although the sequence space of domain-sized fragments appears intractable due to its size, protein scientists have nevertheless developed expectations about its use in nature through decades of biological and bioinformatic research. This is because most proteins have arisen by descent and differentiation from a set of prototypes and can thus be classified into a hierarchy of approximately 10^5^ families and 10^4^ superfamilies (8–10), which represent a negligible part of the available sequence space. This points to the fact that the space around proteins is substantially populated by their homologs, resulting in an image of local islands of natural sequences within a global sea of virtual, unrealized possibilities (11–13). This view is supported by the strong constraints on sequence conservation exerted by divergent evolution. For example, the ubiquitin of humans and budding yeast is still 96% identical despite at least 2 billion years of cumulative evolutionary distance and, while this is certainly an extreme case, considerable sequence identity can be observed in other important proteins of the cell, such as citrate synthase or the core complement of ribosomal proteins, many having diverged at the time of the last universal common ancestor, 3.5 billion years ago. The constraints of divergent evolution still manifest themselves, albeit to a reduced extent, even in proteins that have lost the functionality common to their parent family, such as P-loop NTPases that have lost the ability to bind nucleotides and therefore show a decayed P-loop motif, yet are readily identifiable as homologs by common sequence comparison methods. There is thus a structure of natural protein sequence space which appears obvious to biologists, with some of the islands they envisage assuming great sizes of hundreds of thousands of members, for example in the dominant families of metabolic enzymes, Rossmannoid, flavodoxin-like, P-loops and TIM-barrels, all belonging to the *α*/*β* structure class. These families show further sequence similarities, which have been variously interpreted as the remnants of an ancestral common origin (14), suggesting that they are themselves not positioned randomly in sequence space relative to each other The clustered nature of natural proteins is thus strongly supported by many biological observations and is in fact the predominant view among biologists (15–17). The extent to which this view of islands in an ocean of virtual sequences approximates the structure of global sequence space is however unclear.

In a contrasting view, information theoretical analyses found that the entropy of natural sequences due to sequence correlations is only negligibly lower than the entropy of random sequences (24, 25). Further, considerations of biophysical constraints acting on protein structure also suggested that natural sequences are only ‘slightly edited’ versions of random sequences (18). Searches for differences between natural and random sequences yielded several features such as hydrophobicity patterns (19–21) and preferences for secondary structures (22), but their relevance has remained controversial. A comparison of length distributions for consecutive hydrophobic and hydrophilic stretches returned comparable values for sequences judged to be structured on a 3D lattice model and for random sequences(23), suggesting that foldable protein sequences cannot be distinguished from random ones by these patterns alone. Related studies attempting to find the effects of homologous descent and secondary structure formation on the frequency of natural pentapeptides showed that the frequency of some deviated substantially from random expectation, but that, overall, the frequencies were close to random(5, 6).

The conclusion that natural sequences are not easily distinguishable from random ones, however, did not really reach the biological community and the view of protein families as islands in sequence space remains pervasive. Reasons for this may include a perceived lack of significance of simplified representations for the properties of natural proteins (for example the use of lattice models (23) or pentapeptides (5, 6)) and a description of the findings in terms not readily accessible to a biological audience, but ultimately, it was the inability of these studies to account for the structuring of sequence space observed in biological studies that led to the perpetuation of opposing views in the two communities.

To evaluate these opposing views, we used methods familiar to both communities in order to analyze protein sequences of biologically relevant length, derived without simplifying assumptions from the genomic data of more than a thousand bacterial species, chosen for phylogenetic breadth and controlled for redundancy. We compared this natural dataset to several random ones generated by established random sequence models that incorporate the compositional biases of natural proteins at different levels. We carried out the comparisons with a distance-based approach, which can capture features of how sequences are distributed in global sequence space. Studying the global distribution of data points in space through pairwise distances is common in other fields, such as protein structure determination (26), spatial statistics (27) and economics (28), but has not, to our knowledge, been applied to investigate the global distribution of protein sequences. Specifically, we derived pairwise distances between fragments of a given length within each dataset as the inverse similarity scores of their pairwise alignments. The resulting probability mass function of distances within the set of natural or random sequences are then compared to each other. This approach allowed us to study sequences of up to 100 residues in length, approximating domain size and far longer than any whose global distribution has been studied to date.

We find that, contrary to our expectations and in support of the views on global sequence space derived from biophysics and information theory, around 99% of all distances in natural proteins can be modeled by a random dataset incorporating the natural amino acid composition at the level of individual proteins. We further estimate that of the remaining 1% of distances most are due to convergent sequence features, leaving the proportion of distances unambiguously assignable to homology in the range of 0.1% - clearly detectable but much smaller than anticipated. It is astonishing that this slight deviation is sufficient to account for the observed structuring of the natural proteome into families, superfamilies, and folds.

## Results

### Choice of a natural dataset

For an adequate dataset that reflects the natural protein sequence space, we aimed to achieve a reasonable coverage of deep phylogenetic branches with complete and well-annotated proteomes. Given that the genome coverage for the archaeal and eukaryotic lineages is still sparser than for bacteria, and that particularly eukaryotic genomes are affected by issues of assembly, gene detection, and intron-exon boundaries, we built our database from the derived bacterial proteomes collected in UniProt (29).

To control for redundancy, we selected only one genome per genus and filtered each for identical open reading frames and low-complexity regions. In total, our dataset comprises 1,307 genomes, 4.7 · 10^6^ proteins, and 1.2 · 10^9^ residues. We simplified complexity arising from the use of modified versions of the 20 proteinogenic amino acids, which occurred in a few hundred cases, by converting these to their unmodified precursors, thus maintaining an alphabet of 20 characters throughout. Further details on the generation of our dataset and its specific content are provided in the methods section and supplemental information. In order to contrast natural similarities with randomly expected similarities, we developed a series of increasingly specific random models that account for compositional effects.

### Random sequence models

Our most basal model considers completely random sequences of the 20 proteinogenic amino acids, in which each occurs with an equal probability of 5% (E-model). This model is known to approximate natural sequences only poorly. This is hardly surprising as natural amino acid frequencies in fact range between 1% and 10%, a bias which is associated with metabolic pathways, bio-availability, and codon frequency. We therefore built models that factor in this compositional bias at increasingly local levels.

The first model incorporates the global amino acid composition of our natural dataset, which we refer to as the A-model. More specific models consider local fluctuations in composition. The composition of different genomes, for example, varies with mutational biases, GC-content (30) and environmental influences (31, 32). This effect can be factored in using the individual genome composition (G-model). With a more local focus, compositional bias can also be accounted for at the level of individual proteins (P-model) (33, 34) or even at the level of their fragments (F-model). Having accounted for all compositional effects, the remaining differences between natural and random sequences of the F-model must be attributed to sequence effects, due either to divergence from a common ancestor or convergence as a result of similar evolutionary constraints such as secondary structure formation.

### Sequence similarities and their distribution

Sequence space has frequently been analyzed with a direct approach based on the exhaustive enumeration of natural k-mers, and the comparison of their frequencies to those derived from a random model (5, 6). This approach is restricted to k-mers of length five or smaller, due to sequence space complexity and the data sparsity caused thereby. Furthermore, focusing on abundance of peptides reflects the most *local* approach to studying the occupation of sequence space. An increased density of natural sequences in the nearby neighborhood, for example of point-mutation neighbors, is not considered, as studied in (35–37). An approach based on exhaustive enumeration disregards the relative position among k-mers, which would result in a *global* perspective of how sequences are distributed in sequence space.

We use an indirect approach to circumvent the issues of data sparsity and local consideration of sequence space occupation. Our approach is built on the probability mass function of pairwise distances between sequences of the same length, in the following referred to as *distance distribution*. The distance distribution illustrates how often two strings (in our case protein sequences) are positioned at a certain distance relative to each other within sequence space. This representation allows to capture aspects about the global distribution of considered sequences, which we will detail in the following sections. By comparing distance distributions it is also possible to study differences between sets of sequences. Some distances between natural sequences may be small and unlikely (relative to a statistical background model) and others may be more likely. However, it is the *frequency*, also of unlikely distances, that can give clues about the global distribution of sequences in sequence space. By using lengths of up to 100 residues, our sequences reach domain size (7).

Clearly, a crucial aspect of our approach is the definition of distance in sequence space. Here, distances are defined by normalizing alignment scores and converting them to distances (see methods section). Our main conclusions do not depend on the exact distance used. In detail, we varied all standard parameters of alignments, which are match/mismatch scoring, gap penalties and mode (Smith-Waterman for local or Needleman-Wunsch for global alignment). We present the results of a Smith-Waterman alignment with identity matrix scoring throughout the paper and provide results from other metrics in supplemental information.

The reasons why we use the identity matrix are twofold: (I) as we aim to extract differences between natural and random sequences, using similarity matrices like PAM or BLOSUM induces an evolutionary or biophysical bias to our method. (II) similarity matrices like PAM or BLOSUM do not define distances in a metric space (see supplemental information) as they do not satisfy positivity and the triangle inequality requirements. We note, however, that the mPAM matrix (38) which satisfies these metric constraints and also includes information about the biochemical similarity of amino acids achieves similar results to the identity matrix.

In this context, it is important to note that our method differs from common homology-based approaches. In many cases, evolutionary approaches compare sequences and further iterate comparisons in order to extract patterns of conserved residues, as indicators of homologous relationships. Thereby, these approaches focus on representing evolutionary relationships, visualized as islands in sequence cluster maps. Our distance-based approach is not designed to extract such relationships between remote relatives. Instead, we primarily aim for an unbiased view onto sequence space, that can describe how natural sequences are distributed therein. We expect that differences between the distribution of natural and random sequences truly reflect the impact of natural biases on their distribution in sequence space.

Studying the global distribution of data points in space through pairwise distances is common in other fields, such as protein structure determination (26), spatial statistics (27) and economics (28), but has not, to our knowledge, been applied to investigate the global distribution of protein sequences. Such distance-based approaches do not preserve information about specific positions of data points in space, but rather characterize their *global clustering* and *dispersion*. Similarities between different types of sets, even if they do not share any members, can be compared by their distance distribution.

Comparing a set of natural sequences to a set of sequences generated from one of the random models, can then be reduced to subtracting the two resulting distance distributions. We refer to this difference as the *residual*. Over all alignment scores, residuals sum up to zero and may locally have values that are either positive (more natural distances) or negative (more random distances). In order to obtain an overall measure of how different two distance distributions are, we derive the *total residual*, which is the variational distance between two probability mass functions.

More precisely, the total residual is the sum over the absolute residuals, normalized to a range between 0% and 100%. If the two distance distributions are completely non-overlapping, the total residual assumes the maximal value of 100%, indicating that no distance between natural fragments can be modeled with random sequences. If they are identical, the total residual assumes a value of 0%, indicating that 100% of distances in the natural distribution have a corresponding distance in the random distribution. Thereby, the total residual represents the fraction of natural distances that are not accounted for by the distance distribution of a random model. We note, that we also evaluated our results using the Kullback-Leibler divergence. This metric leads to similar results and are further presented and discussed in the supplemental information.

### Global amino acid composition bias

We start our analysis by assessing to what extent the global amino acid composition, as captured in the A-model, can describe natural sequences. We compare the distance distributions of the two datasets for fragment lengths of 10 up to 100 residues, in increments of 10. At all fragment lengths, the results are closely comparable. We show the results for 100-mers as representative for domain-sized sequences in (Figure 1) and provide the others in the supplemental information.

**Fig. 1.**
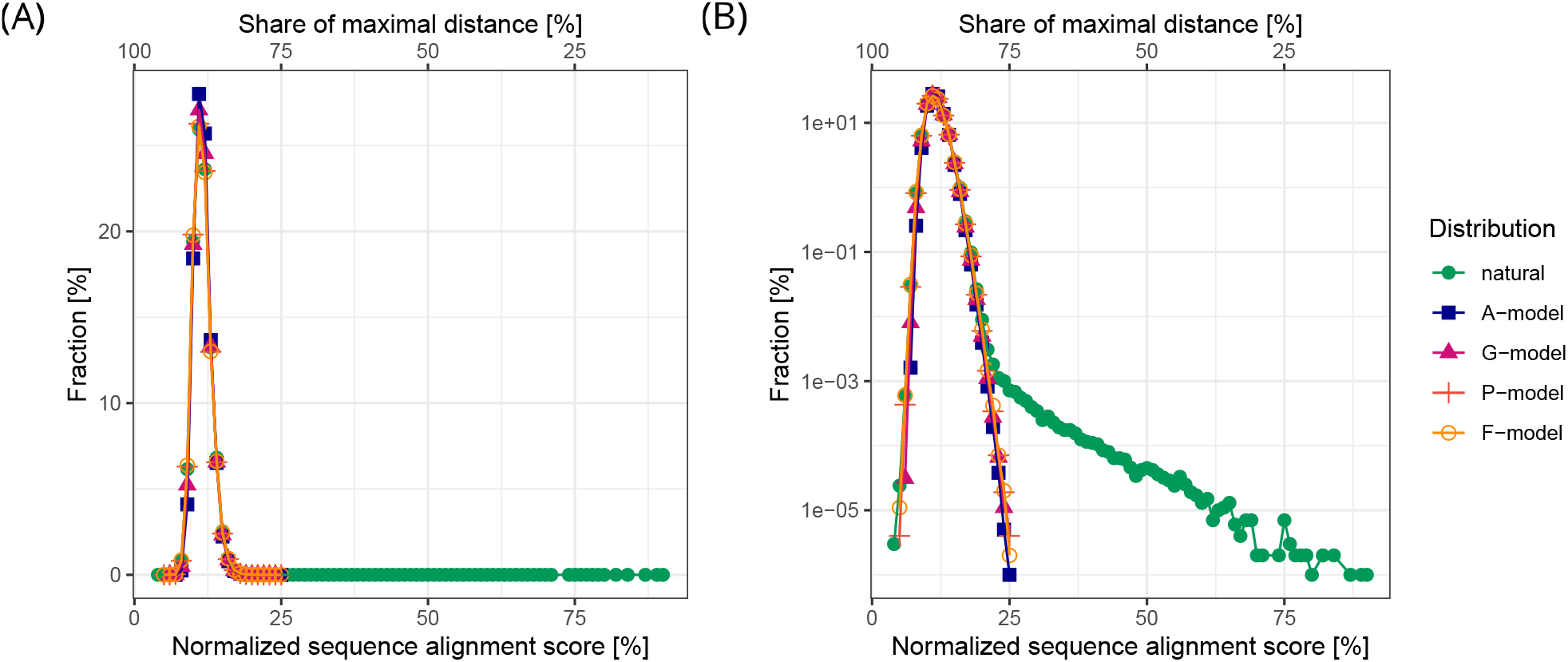
Sequence similarities of random protein sequence models and natural sequence data. (A) Distance distributions as a descriptor of how natural sequences are distributed over sequence space. The distances between sequence fragments of the same length are defined by the sequence identity score of a Smith-Waterman alignment. The scores of 100 Million pairs are plotted as a probability mass function (raw data is provided in the supplemental information). Values of zero were omitted. All distance distributions spike in the area of long-range distances with a mean sequence identity score around 11%. Both natural and random distance distributions are almost entirely overlapping. (B) Same data with a log-scaled y-axis. With this representation, a small fraction of significant similarities between natural fragments becomes obvious.

The distance distributions of natural data and A-model overlap extensively (Figure 1: A). Both are uni-modal with a peak at a low alignment score of 11%. Despite their considerable similarity, however, they differ in two respects. (I) Natural sequences have a small but measurable fraction (below 0.001%) of distances closer than what amounts to 25% sequence similarity, whereas the A-model has none (visible only in a log-scale plot of the distance distributions, Figure 1: B). We will show further on that these distances correspond to homologous relationships and note that the region around 25% pairwise sequence identity has been previously described as the “twilight zone", below which homology cannot be reliably inferred (39, 40). (II) The larger difference between the distance distributions occurs below scores of 25% sequence identity and is best visualized by their residuals (Figure 2). These take the shape of a wave, with two maxima at normalized similarities of 9% and 15% (reflecting an over-representation of the corresponding distances in the natural dataset), and a minimum at 11% (reflecting an under-representation). The over-representation of distances both longer and shorter than expected from the random model, suggests that natural sequences are less homogeneously distributed in space.

**Fig. 2.**
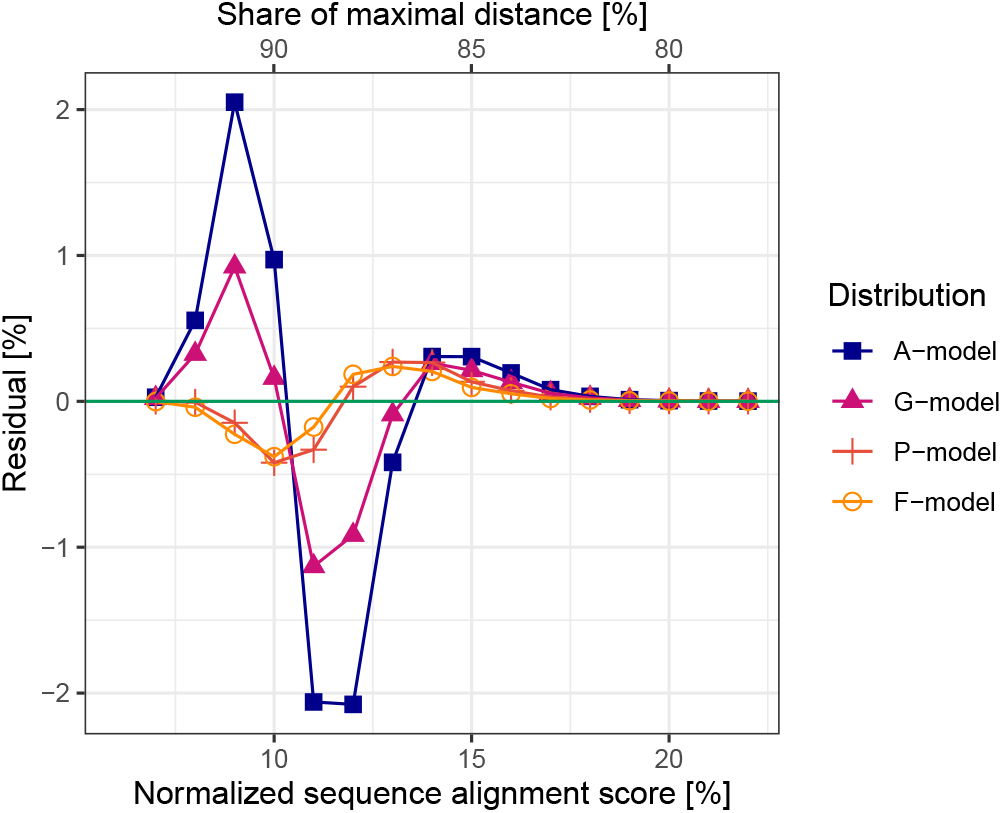
Difference between the distance distributions of natural sequences and random models. We extract the distance-specific difference by subtracting the random from the natural distance distribution. The resulting residuals for each model indicate distances between natural fragments that are unaccounted for by the respective model (crests above zero). The A- and G-model display a 2-peak behavior, associated with both more long-range and short-range distances between natural fragments than modeled, reflecting an increased amount of both diversity and clustering in natural sequence space. The residuals of the P- and F-model possess only one peak for more short-range distances between natural sequences, hence an unexpected amount of clustering.

We rationalize this effect with the observation that natural polypeptide chains tend to bias their composition towards a subset of available amino acids, thereby showing a reduced entropy relative to random sequences of the A-model (Figure 3). This may include a bias towards rare amino acids (such as Cys, Trp and His in small proteins induced by the biological necessity for zinc-coordination and disulfide bridges (41)) as well as towards abundant amino acids (such as Leu, Ala and Glu in all-alpha proteins, most extremely in coiled coils (42)). These compositional biases mean that distances in sequence space between proteins of the same bias will be shorter than expected from the A-model, while they will be longer between sequences showing different biases. Since residuals add up to zero, the number of intermediate distances is correspondingly decreased.

**Fig. 3.**
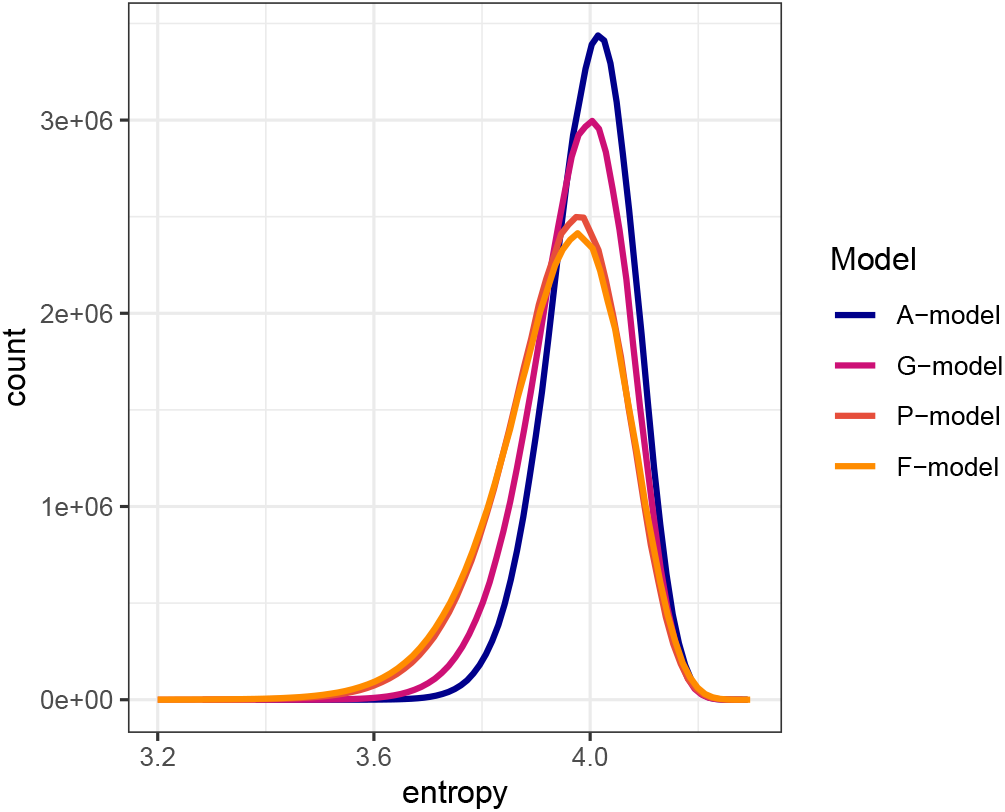
Compositional entropy of 100-mers from different random models. Fragments from the A-model possess the highest entropy on average. With more local consideration of the natural composition, the entropy reduces. Fragments of the F-model possess the same entropy as the natural 100-mers.

We note, however, that this discrepancy between natural sequences and the A-model is not very pronounced, as the total residual has a value of only 4.5% for 100-mers (Figure 4). It is even less pronounced at smaller fragment lengths, reaching 0.4% for 10-mers. Thus, the A-model becomes less accurate in describing the sequence space occupation of natural sequences at lengths that are biologically relevant, but it already achieves considerably higher accuracy than the completely random model (E-model), which has a total residual of 30.4% for 100-mers (supplemental information).

**Fig. 4.**
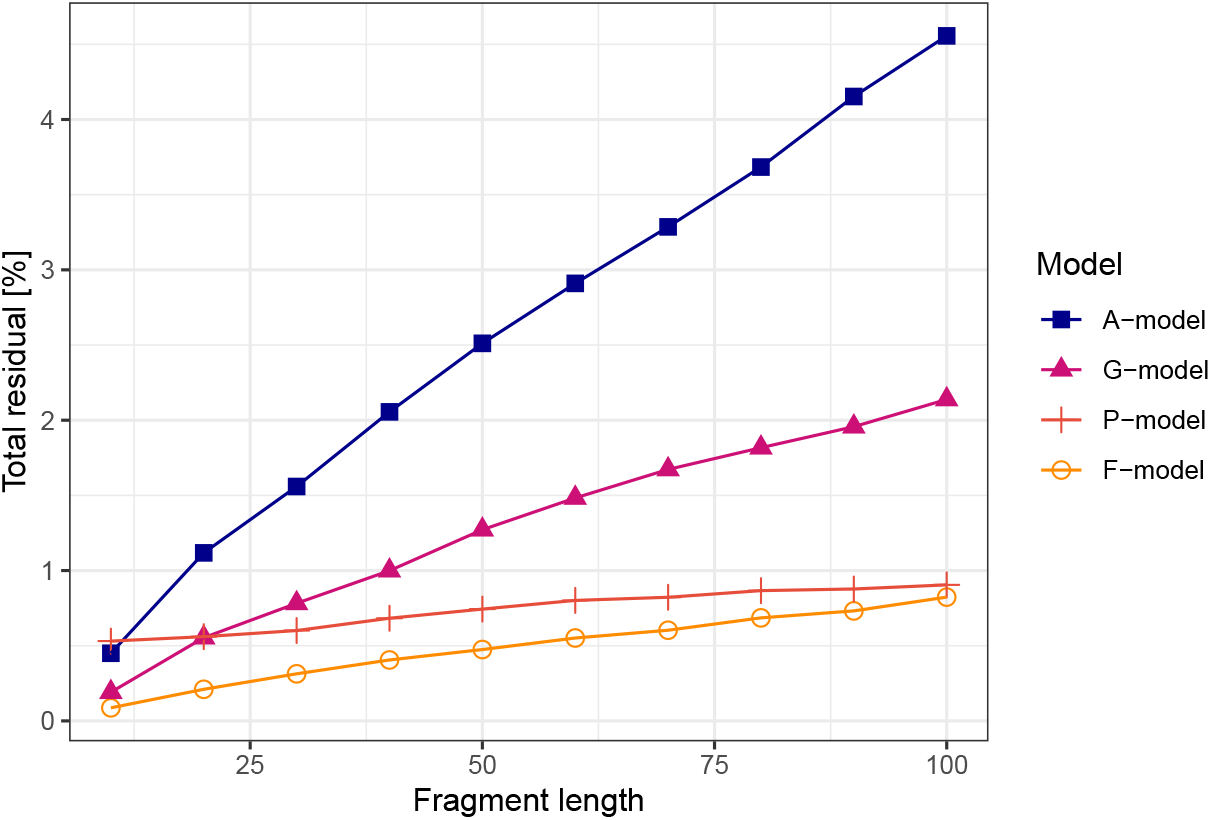
Deviation of random sequence models from natural sequences as a function of fragment length. Total residuals when using a local Smith-Waterman alignment The total residual indicates the extent to which the distance distribution of random sequence models deviates from the natural. It reflects the fraction of distances between natural fragments that are unaccounted for in the random model. With increasing fragment length, the total residual of all models increases, implying that for longer fragments all models become worse in approximating similarities between natural fragments. The A-model (overall amino acid composition) deviates furthest followed by the G-model (residue composition of genomes), the P-model (residue composition of proteins) and the F-model (residue composition of considered fragments), which deviates the least. The intercept of the total residuals of the P-model with the other models at fragment length around 10 is associated with edge effects of natural sequences and the usage of a local alignment as distance metric.

We conclude that a random sequence model based on the amino acid composition of the considered proteins already approximates the distribution of natural sequences over the global sequence space fairly accurately and we expect further improvement by including compositional biases on a more local level.

### Context-specific amino acid composition

As a first step we considered a model that accounts for genome diversity (G-model). Therein, the random dataset is produced by shuffling residues of the natural dataset within the boundaries of each genome. Given that our natural dataset holds 1,307 genomes, the derived sequences are thus sampled from 1,307 distinct compositions, which introduces the compositional heterogeneity of different genomes into the random dataset (Figure 3). Comparing the G-model to the A-model over the bacterial dataset shows a dampened wave for the residuals, with the same shape, but a decreased amplitude (Figure 2). The total residual for the G-model is thus only about half as large as that of the A-model (Figure 4), implying that controlling for genome composition provides a substantial improvement in modeling the natural distance distribution.

Further locality is achieved by accounting for the composition of individual proteins (P-model). Here, the random dataset is produced by shuffling residues within each natural protein, corresponding to 4.7 · 10^6^ compositions. This model yields a further improvement over the G-model, even though, at sequence lengths below 20 residues, it produces minor inconsistencies in its total residuals relative to the other models (Figure 4). We suspect that this is an artifact of using local alignments and, indeed, the effect disappears when using a global alignment as distance metric over the same dataset (see supplemental information).

The shape of the residuals of the P-model is qualitatively different from the A- and G-models. It has only one maximum near a normalized similarity of 13%. The maximum at smaller similarities, that is evident in the A- and G-models, can no longer be observed which we attribute to the fact that accounting for composition at the level of individual proteins has introduced almost all of their their compositional bias into the random model (Figure 3).

For 100-mers the total residual of the P-model is 0.9% (Figure 4), a value that is not improved remarkably by an even greater locality at the level of individual k-mers, yielding a total residual of 0.8% for the F-model.

### Decomposition into homologous and analogous relationships

Having accounted for compositional effects at increasingly local levels, the remaining discrepancy between the distance distributions of the F-model and the natural dataset must be due to the actual sequence of amino acids, such as specific k-mer biases. This discrepancy can arise either through divergence from a common ancestor (homology) or convergence as a result of structural constraints, particularly secondary structure formation (analogy). In order to evaluate the relative contribution of these mechanisms to the natural distances between sequence fragments we aimed to identify what proportion of distances could be assigned confidently to either homologous or analogous relationships.

The detection of homologous relationships requires advanced approaches, which are computationally much more expensive than the simple sequence alignments used to determine distances in sequence space. In order to render this problem tractable, we sampled a subset of our sequences, which was unbiased and large enough to accurately reproduce the total residuals in the previous analysis (0.82% in both cases, compare Figure 4 and Figure 5: D). For this, we assembled 10 unbiased groups of 100-mers from our natural dataset, containing approximately 650 sequences each. We used HHblits (43) to generate profile Hidden Markov Models (HMMs) for all individual sequences within these groups, then sampled relationships between sequences belonging to different groups by aligning their HMMs in HHalign. This resulted in multiple unbiased samples of relationships, which amounted to 4.6 million fragment pairs.

**Fig. 5.**
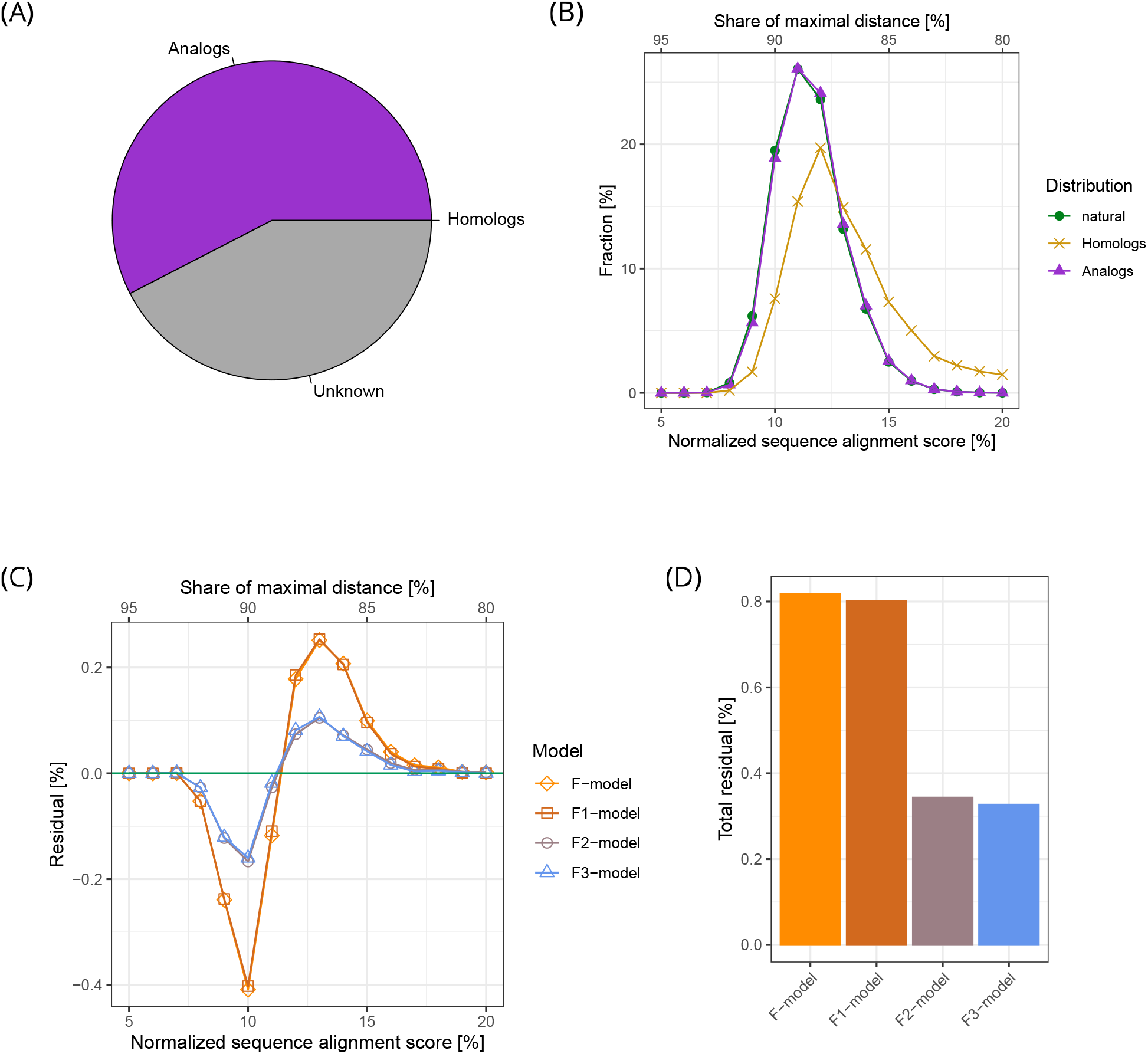
The contribution of homology and analogy to the global usage of sequence space. (A) Decomposition of fragment pairs into their origins. We sampled 2 Million fragments pairs and analyzed if their relationship is confidently homologous or analogous. The fraction of analogous relationships was determined to be 57%, homologous relationships only 0.1% and the remaining fraction of is of unknown origin. (B) Distance distribution between homologs and analogs contrasted with the natural distance distribution. The qualitative difference between the distance distribution of analogs and that of all fragments is relatively small. Compared to this, the distance distribution of homologs displays a strong tendency towards a higher sequence identity score; it nevertheless has a major overlap with the natural distribution. (C) Residuals of the models incorporating the sequence bias of homology and analogy. We generated mixed models, that include the sequence bias of homology (F1-model), analogy (F2-model) and both (F3-model) into the F-model, which is only based on the composition of natural 100-mers. The F1-model, which includes homologous sequence bias, displays almost the same residuals as the purely composition-based F-model. The residuals of the F2-model, which includes analogous sequence bias, deviates severely from that of the F-model. The F3-model yields similar results as the F2-model. (D) Total residuals of mixed models of 100-mers. The total residuals behave according to the residuals. The total residual of F-model over the sampled fragment pairs is 0.82%, which is in accordance with the total residual of the F-model over the entire dataset. The F1-model displays an only minor improvement in the total residual of 0.016% compared to that of the F-model. The F2-model reaches a total residual of 0.34% and is more than 2-fold more accurate than the F-model. Adding the homology bias to the F2-model to obtain the F3-model has almost no effect.

We assume fragments are confidently homologous, when their derived HMMs have a minimal probability of 90% to be homologs according to HHalign (see methods section). This process identified more than 4,900 distances to be homologous, which corresponds to 0.10% of the total sampled distances (Figure 5: A, yellow). The standard error of the mean (SEM) between the 10 sampled relationships was 0.0039% (Figure 5: A). More detailed numbers can be found in the supplemental information.

For the remaining sequence pairs, we evaluated the likelihood of analogy by comparing their HMMs to those of the ECOD database (8). By virtue of containing only domains of known structure, ECOD is the currently best resource for distinguishing between homology and analogy in protein domains. For our analysis, we classified pairs of sequences as analogous if they matched distinct X-groups in the ECOD hierarchy. The matching of fragments to a X-group was facilitated by aligning the fragment’s HMM against HMMs generated for each X-group via HHalign. The match was accepted when HHalign predicted a probability of homology above 90%, which is the same cutoff used for the homology assignment between fragments. In most cases, the X-level is the highest level at which homology still needs to be considered as a possibility; requiring fragments to match different X-groups thus provided a conservative estimate of analogous relationships. This process identified 57% of pairwise relationships as analogous (Figure 5: A, purple), with a SEM of 0.63%. The remaining 42.8%, with a SEM of 0.63% (Figure 5: A, grey), comprise sequence pairs which we could not assign to either group.

If we use a less stringent cutoff of 50% probability the proportion of unknown relationships decreases, while the ratio between analogous and homologous relationships remains similar. With this decreased cutoff we can assign 0.19% of relationships to the homologous and 72% to the analogous group (see supplemental information). The remaining relationships could not be determined due to the lack of assigned structure to either of the considered fragments and are thus thought to contain mostly relationships between unstructured or non-canonical regions.

We conclude from this that the number of analogous pairs exceeds the number of homologous pairs by more than two orders of magnitude. This indicates that the influence of homology on the global distance distribution in natural sequences is dwarfed by analogy.

### Sequence bias caused by homology

Having decomposed sequence pairs into homologous and analogous relationships, we analyzed to what extent the remaining total residual of the F-model can be explained by incorporating corresponding sequence biases into our F-model. Therefore, we generated three new hybrid models in the following way: we omitted either homologous pairs, or analogous pairs, or both from our set of assigned relationships, generated an F-model for the remaining fragment pairs through the same shuffling procedure as used previously, and then added back the omitted pairs without shuffling. In the following we refer to the hybrid model that adds the sequence bias of homologs to the fragment composition as the F1-model, the one that adds the sequence bias of analogs as the F2-model, and the one that adds both biases as the F3-model.

Relative to the total residual of the purely compositional F-model, the F1-model, which includes homologous sequence effects, is only minimally more accurate (total residual reduced by 0.016%) at approximating the natural distance distribution. Two reasons are mainly responsible for this only minor improvement: First, the proportion of homologous relationships is only 0.10%, giving them little leverage. Second, the distance distribution of homologs differs only to a small extent from the distance distribution of the natural dataset (Figure 5: B). In fact, it has been recognized previously that most homologous sequences share no significant similarity (44, 45). This observation indicates a broad distribution of homologs over the entire sequence space. Clusters of homologous sequences, especially those with ancient relationships, are thus diffusely distributed and not compactly located at a location in sequence space.

In contrast, the total residual of the F2-model, which includes analogous sequence effects, is decreased about 2.4-fold relative to the F-model (from 0.82% to 0.34%), as visualized in Figure 5: D. Thus, although analogs have a distance distribution that is very similar to the natural one, their leverage is 2 orders of magnitude higher than that of homologs, causing these small differences to improve substantially the fit of the F2-model to the natural distance distribution. As the compared sequences were assigned to ECOD domains, they share the ability to form secondary structures, resulting in a sequence bias that is not captured by residue composition. Adding the homologous sequence bias to the F2-model to generate the F3-model did not improve its ability to approximate the natural distance distribution.

We conclude that although divergent evolution is the main mechanism populating sequence space, its impact on the global distribution of sequences is only minor. Instead, compositional effects shape the global use of sequence space almost entirely and the marginal impact of sequence biases is mostly due to analogy, which might result from the required ability to form secondary structure.

## Discussion

In this article we have undertaken a study of natural protein sequence space, using an approach built on the probability mass function of pairwise distances between sequence fragments. With this approach we were able to analyze the occupancy of sequence space by fragments up to 100 residues in length, substantially going beyond previous efforts and for the first time characterizing their global distribution in space. Our results show that the compositional bias of natural proteins is already sufficient to approximate the distance distribution of natural 100-mers by more than 95% and that accounting for local compositional bias down to the level of individual proteins or fragments further improves this to more than 99%. The distances that remain unaccounted for are mainly contributed by sequence effects arising from analogous relationships, leaving only a minor contribution to homology in the global characterization of sequence space occupancy - clearly detectable but much smaller than expected..

This surprised us, as decades of bioinformatic work have mapped out an increasingly comprehensive description of sequence space around protein families, based on the detection of ever more remote homology. Clearly, local sequence space is dominated by the legacy of evolutionary history, as seen for example from the fact that at sequence identities above 30%, the space around a protein is populated almost entirely by its homologs (39). We therefore expected to find that homology also has a substantial role in shaping the global structure of sequence space, corresponding to the image of islands formed by natural sequences within a global sea of possibilities. This expectation was not borne out and in retrospect this might not seem as surprising, given that even within protein families, the influence of homology is smaller than generally perceived. This is due substantially to the way in which family relationships are represented, strongly emphasizing common features (such as the generally few highly conserved residues (44)) and omitting variable ones. This focus on biological significance over raw sequence similarity leads to a perception of sequence space that is dominated by evolutionary connections, not by the pairwise sequence identity. Evidence for this can be seen for example in the progressively more complex statistical methods needed to substantiate homology across increasingly large evolutionary distances, the resulting difficulties to classify the detected relationships into a hierarchy of protein families and superfamilies, and the remaining inability in many cases to judge on the homologous or analogous nature of similarities, even in the presence of extensive sequence and structure information. These considerations show why, even at the level of protein families, many sequence relationships share only residual similarity below 20% identity, which is fundamentally indistinguishable from that arising by random drift or convergence. Thus, if we denote the region of 20–35% pair-wise sequence identity, where the proportion of analogs grows rapidly, as the ‘twilight zone’ and the region below it as the ‘midnight zone’, most homologs of a protein are found in the midnight zone (39).

We conclude that at sequence relationships below 40% identity the image of islands, which is a good descriptor of local sequence space, rapidly breaks down. Whereas islands grow in two dimensions, sequence space grows in twenty. This means that, already at a fragment length of 62 amino acids, the number of possible sequences exceeds the number of particles in the known universe. In this space, whose occupation becomes extraordinarily sparse at low sequence identities, analogous and random similarities rapidly overwhelm homologous ones in number. We therefore think that the image of clouds with ever more diffuse edges is more suitable to describe homologous families in global sequence space.

These considerations provide a framework for understanding why it was so difficult to identify a global impact of homology on the distribution of natural sequences in space. Rather, we conclude from our results that neither divergent evolution from a common ancestor nor convergent evolution due to biophysical constraints prevent the exploration of global sequence space by proteins, although they may slow down the process dramatically due to functional requirements. In this view, the extreme sparseness of proteins in sequence space is mainly due to its gigantic size and to the overwhelming proportion of non-functional forms, since in biological evolution all trajectories connecting two proteins must pass exclusively through functional intermediates. Thus, while there may be many foldable and potentially functional forms located far from currently observed proteins, these may not be reachable by natural evolution and only become accessible through the arsenal of new methods in protein engineering and design.

## Materials and Methods

### Details to the used natural data

We selected the majority of bacterial genomes provided by UniRef on 22.09.2017 (29). Some genomes stood out as they possessed multiple replicas of the same protein and were excluded, leaving 4,097. For each of the 1,307 genera we randomly chose one representative for our natural dataset. The considered and selected genomes are provided in the supplemental information. The genus was derived from the full-length genome name via string matching. We are aware of the general ambiguity of the definition of a genus (46). However, with the genus selection we only aimed to reduce redundancy caused by some genera or species that have been sequenced many times. Lastly, we note that the bias towards bacteria that are easy to cultivate prohibits a sampling of the true diversity among bacterial genomes.

Apart from redundancy at the genome level, we control for recent gene duplication events. For each genome, we cluster its proteins using cd-hit (version 4.6 with 99% sequence identity and 90% coverage). A representative protein sequence, as defined by cd-hit, was then selected for each cluster; all other proteins were discarded.

Low-complexity regions (LCRs) are a well-known features of natural sequences, that do not occur as frequently in random sequences. We first analyzed our data including LCRs and found that they significantly contribute to the total residual between natural sequences and our models (data not shown). Therefore, we pruned LCRs of our natural dataset using segmasker (47) (version 2.3.0+ with the standard settings), to study differences between natural and random sequences that are not due to this well-known feature. This pruning of LCRs leads to natural sequences of slightly higher complexity than that of the random sequences (data not shown). This bias plays an insignificant role, especially for longer sequences, which are of most interest in our study. Since N-terminal methionines were sometimes included, we stripped them to standardize our sequences.

To simplify our analysis we changed a couple of hundred cases of uncommon amino acids to their most similar proteinogenic amino acid. In order to use the exact same dataset for all sequence lengths, we pruned our dataset of sequences shorter than 100. Additionally, we removed the invalid amino acid X by replacing it with an end-of-line-character, effectively dividing a protein sequence into multiple parts. However, since some of our random models depend on shuffling intact genomes or proteins, we performed this division into multiple parts after the shuffling (more detail below).

Taken together our dataset holds 1.2 · 10^9^ valid amino acids of 1,307 genomes comprising 4.7 · 10^6^ proteins.

### Data and code availability

The code for generating the models and datasets used in this study is available under the GLPv3 license. Code and documentation are available in a GitHub repository at https://github.com/Lolasdasdasd/Sequence_Space. We also provide other additional data in this repository such as the raw data of the derived distances, a list of the considered and used genomes.

The natural dataset is available at ftp://ftp.tuebingen.mpg.de/pub/protevo/lweidmann.

### Fragment pair selection

In the process of generating distance distributions, pairs of fragments with identical lengths need to be selected. We selected pairs of fragments in an unbiased and randomized manner such that each character (amino acids and end-of-line-character) in the dataset had the same probability of being chosen as the initial letter of fragments. Fragments were then extracted by extending the string starting at the chosen initial letter to the respective length. Those fragments that straddled protein boundaries or invalid regions, indicated by the end-of-line-character, were rejected. Our random selection of fragment pairs was designed to have a cycle length much larger than the number of fragment pairs used for calculation of a distance distribution. We could therefore avoid to chose the same pair twice. We ensured this by implementing two linear congruential generators (48) to enumerate all possible pairs of fragments. In detail, one linear congruential generator was used for each member of the pair with multiplier *a* = 1 and moduli *m*_1_ = 223 and *m*_2_ = 34, 211, where both moduli are prime numbers relative to the total number of characters 1, 168, 754, 000. Depending on the starting points of the two generators, a different subset of index pairs will be selected. This enabled us to calculate disjunctive fragment pairs in parallel. We selected 10^8^ valid pairs of fragments to accurately estimate the distance distributions.

### Randomization in all random sequence models

All our random models are based on random permutations, for which we used the Mersenne Twister algorithm mt19337 of the C++ 14 std library with the standard seed value of 19650218. This algorithm is considered one of the best pseudo-random number generators and in a test with a smaller dataset we found that our results did not depend on the type or seeding of the random number generator. For the E-, A-, G-, and P-models, we generated permuted datasets (using C++ function random_shuffle) of the same size as the natural data and selected fragment pairs with the algorithm described above. For the F-model we did not generate any additional dataset. Instead, we generated permuted versions of the considered natural fragments in the process of fragment pair selection for the calculation of the natural distance distribution. For the F1-, F2-, and F3-models a limited number of fragment pairs was sampled from the natural dataset. This required fragment pairs to be chosen differently from the other models to ensure an unbiased sample. Of all the selected pairs, the distance between the original, natural fragments and that between one of their permuted versions (generated by the unix command shuf) was derived. Pairs were further classified into homologous, analogous and unknown relationships and, depending on the desired bias, the distance between the original or permuted fragments was reassembled. All models are explained in further detail in the following.

### Models incorporating overall amino acid composition

The first random sequence model, which we refer to as the A-model, is based on the amino acid composition of the whole natural dataset. We obtained randomized sequences for this model by randomly shuffling all amino acids of the natural data. Thereby, protein length is maintained and the number of amino acids stays exactly the same. We note, however, that the specific propensity of amino acids at positions near the ends of natural sequences is not reflected in our random model.

For the E-model, we proceeded the same way as for the A-model. The only difference is that we replaced the natural dataset, by writing over all valid amino acids with the 20 possible amino acids in lexicographical order.

### Models based on the amino acid composition of individual genomes or proteins

To account for the composition of individual genomes or proteins, we permuted amino acids within the context of genomes or proteins. For the G-model, we shuffled valid amino acids within each of the 1,307 genomes. For the P-model we shuffled valid amino acids within each protein. We used one instance for genome and protein composition bias and stored them to generate the distance distribution for the corresponding models. Multiple instances for genome and protein composition bias were found to converge to the same results after a large enough sampling of more than 10^8^ fragment pairs. After shuffling, we divided proteins containing the invalid amino acid X by replacing it with an end-of-line-character.

### Model based on the amino acid composition of fragments

For the F-model, we randomly permuted the considered natural fragments. This procedure is similar to the permutation model used in (5), which was applied to pentamers exclusively and is here used on all considered peptide lengths. In contrast to the previous random models, generating a single randomly shuffled dataset is not computationally feasible since storing an instance of all shuffled fragments would increase the data size approximately by a factor of the respective fragment length. We therefore shuffled fragments on the fly during the calculation of the natural distance distributions. The benefit of this implementation of the F-model is that the exact amino acid composition of the considered natural fragments is captured. Therefore, all deviations must be due to a pure sequence bias. The F1, F2, and F3-models, which incorporate the sequence bias of homologous and analogous fragments, are presented further down.

### Similarity derived form sequence alignments and distance metric

We define the distance between two fragments of the same length *N* as the difference between the normalized rounded similarity score *s* from a Smith-Waterman alignment and the maximal possible score. In the alignment, an amino acid match is scored with 1, a mismatch with 0, gap opening penalty is equal to 3 and gap extension penalty is 0.1, which are the same parameters for gaps as used in (39, 40). We chose local alignments, given that it is the common method to compare natural sequences. In the supplemental informationwe provide results of Needleman-Wunsch alignments and alignments, which lead to similar results. We additionally provide results of different similarity matrices and show that those results depend on the chosen gap open penalty or are inconsistent over fragment length.

Due to gaps, the alignment scores *p* can rank between 0 and *N* in steps of 0.1; to obtain integer distances, we round scores to the closest integer number. Distances exactly between two integers (such as 1.5) are assigned to the smaller one. We note that this binning procedure is done to merge scores effected by gap insertion with the most frequent ones, where no gap was inserted. We chose the smallest binning granularity that avoids this artifact. In principle, bins could also be chosen larger, which however would reduce the resolution of the results. At the chosen resolution leads to the same results when no binning was applied (see supplemental information). To compare the similarity score *p* across different fragment lengths *N*, we transform it into the normalized score *s*, ranging between 0 and 1, as follows:

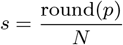

This score *s* thereby reflects the number of dimensions sequence space (positions in sequence), which are the same between two fragments, while allowing for gaps and insertions. In order to obtain a measure of distance *d*, we subtract the normalized alignment score from the maximal possible normalized score, which is 1:

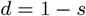

The values for both similarity *s* and distance *d* range between 0 and 1. We typically represent them in percent in the main text.

We compared results of our distance metric with those calculated with other distance metrics using other alignment methods, including global alignments and different gap penalties. The results are comparable to the results reported in the main text and mostly differ in magnitude. Results with other distance metrics are provided in the Supporting Information. For all alignments, we used the SeqAn C++ library, version 2.4 (49), which enables many sequence comparisons in parallel.

### Distance distribution

Having calculated 10^8^ distances between fragments of the same length, we can count how frequent a certain distance is. This frequency distribution can be represented as a probability mass function. We refer to this function of distance *d* as the distance distribution *D*(*d*). Distance distributions can be derived from the natural dataset *D*_nat_ or from any of the presented random models *D*_rand_.

### Comparing distance distributions with residuals and total residual

We define the residuals as the difference between the natural and random model distance distributions for each possible distance *d*. We use it to demonstrate the qualitative difference between the distance distribution of a random model and that of natural sequences. Denoting the residual by *r*, the random model distance distribution by *D*_rand_ and the natural one by *D*_nat_ we have:

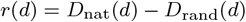

For residuals exceeding zero, there is a higher frequency of these distances between natural fragments relative to that between random fragments. Residuals below zero indicate the opposite, a lower frequency of these distances between natural fragments.

To summarize the difference between natural and random model distance distributions in a single metric, we sum the absolute residuals over all possible distances and normalize it to a range between 0 and 1:

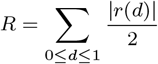

We call *R* the total residual, which is variously called the variational distance, total variation distance or Kolmogorov distance (50). We typically represent the total residual in percent.

### Decomposition into homologous and analogous relationships

We derived the fraction of sequence pairs that are confidently homologous using the tools of HH-suite (version number 3.0.3) (43). To derive this fraction, we systematically sampled our dataset and extracted 10 sets of natural 100-mers that are equally distributed over our dataset, each containing approximately 650 fragments. (See supplemental information for exact sampling sizes.) With HHblits, we generated HMMs with the standard settings for each of these fragments with two iterations, using uniclust30 (51) as underlying database (version August 2018). Then, we pairwise aligned the generated HMMs with HHalign, in order to estimate whether two fragments are homologous. We did this by aligning all fragments in one set to all of those in another set, resulting in 90 possible directed combinations of which we chose 10 as representative sets of pairwise relationships. Each set of fragments was considered twice in this comparison, once as the set of query sequences and once as the set of target sequences in the alignment. This resulted in 2 Million pairwise fragment comparisons divided into 10 disjunctive sets. Pairs of fragments were considered to be homologous, if HHalign predicted them to be homologous with a probability above 90%.

We derived the fraction of sequence pairs that are confidently analogous using a similar procedure as used for the homology detection. We first assigned structured domains to each 100-mer. We then assumed a pair of 100-mers to be of analogous origin, if the two 100-mers matched only distinct domains that are confidently not related to each other.

For the assignment of structured domains, we used the ECOD classification (8), which is currently the best resource for distinguishing between homology and analogy in protein domains. The HMMs of each 100-mer (same as in the homology detection) were thereby compared against all ECOD entries (retrieved on 9.4.2019) with HHsearch. We used HHsearch with the standard parameter and assigned the best-scoring non-overlapping hits with a probability above 90% to the corresponding fragment. Of all 100-mers 70% could be assigned to a single domain and less than 1% to multiple domains, of which we considered both. 100-mers that were not assigned to any domain were directly excluded to be analogous to any other sequence, since we are uncertain about their origin. Pairs containing at least one such fragment without structured domain were classified to be of unknown origin.

For the assignment of analogous relationships, we considered only pairs of 100-mers that were assigned to at least one domain. If all of their domains matched only distinct X-groups in the ECOD hierarchy, the pair was assumed to have an anologous relationship. The X-group is the highest level at which homology still needs to be considered as a possibility. All pairs of fragments that were assigned to domains of only distinct X-levels were considered to be analogous. More details on the derivation of homologous and analogous relationships and the obtained statistical values of the SEM are provided in the supplemental information.

### Mixed models containing sequence bias of homology or analogy

In order to estimate the influence of homology and analogy on the natural distance distribution, we generated mixed models that account for their sequence bias. The F1-model includes the sequence bias between homologs by including the distances between all homologous fragment pairs without shuffling. We applied the F-model to the remaining fragment pairs, that are presumably not homologous. For this, we permuted the fragments of the corresponding pairs with the Unix command shuf followed by deriving their distance. All distances (those of the homologous fragments and those of the permuted, non-homologous fragments) combined resulted into the distance distribution of the F1-model. The sequence bias between homologous fragments is therein preserved while for other fragment pairs only their composition is accounted for, as generally done in the F-model. We proceeded the same way for the F2-model by including the distances of unshuffled fragments that were classified to be analogous, and distances of the remaining pairs after shuffling the residues within each fragment. For the F3-model we included both sequence bias of homologous and analogous natural fragments.

## ACKNOWLEDGMENTS

We thank Mohammad ElGamacy, Joana Pereira, Klara Hlouchova, Chris Sander, Rachel Kolodny and Nir Ben-Tal for insightful discussions and their overall input on this work. This work was supported by institutional funds of the Max Planck Society and by a “Life”-grant from the Volkswagens-tiftung to A.N.L.

